# Evaluation of Methods for Cell Nuclear Structure Analysis from Microscopy Data

**DOI:** 10.1101/254219

**Authors:** Alexandr A. Kalinin, Brian D. Athey, Ivo D. Dinov

## Abstract

Changes in cell nuclear architecture are regulated by complex biological mechanisms that associated with the altered functional properties of a cell. Quantitative analyses of structural alterations of nuclei and their compartments are important for understanding such mechanisms. In this work we present a comparison of approaches for nuclear structure classification, evaluated on 2D per-channel representations from a 3D microscopy imaging dataset by maximum intensity projection. Specifically, we compare direct classification of pixel data from either raw intensity images or binary masks that contain only information about morphology of the object, but not intensity. We evaluate a number of widely used classification algorithms using 2 different cross-validation schemes to assess batch effects. We compare obtained results with the previously reported baselines and discuss novel findings.

## 1 Introduction

Cell nuclear structure is regulated by undelying biological mechanisms related to cell differentiation, development, and disease [3, 10, 11]. Changes in nuclear architecture are related to altered functional properties such as gene regulation and expression. Moreover, studies in mechanobiology show that external geometric constraints and mechanical forces that deform the cell nucleus affect chromatin dynamics and gene and pathway activation [9]. Quantitative analyses of structural alterations of nuclear strucstures also have medical implications, for example, in detection of pathological conditions, such as cancer [11]. Although a few of signal processing and computer vision algorithms have been proposed to analyze cell and nuclear phenotypes using 3D representations [2], the dimensionality of acquired data, various image acquisition conditions, and great variability of cells in a population present numerous challenges for 3D image analysis methods. 2D image representations are computationally cheaper to operate on and do often carry enough information to achieve a desired level of performance.

In this work we present a comparison of approaches for nuclear structure classification, evaluated on 2D per-channel maximum intensity projections from a large 3D microscopy imaging dataset. Specifically, we compare direct classification of pixel data from either raw intensity images or binary masks, which contain only object morphology information, but not texture. We evaluate a number of widely used classification algorithms using 2 different cross-validation schemes to assess batch effects. We demonstrate near-perfect classification performance using 2D data and compare our results with originally reported baselines [5].

## 2 Methods

### 2.1 Dataset description

In this study we use 3D Cell Nuclear Morphology Microscopy Imaging Dataset [5], the biggest public dataset for nuclear structure classification. This dataset contains 3D volumetric microscopic cell images with corresponding nuclear and nucleolar binary masks. It includes images of cells in two phenotypic states that have been shown to exhibit different nuclear structure. Thus, it poses a binary classification problem that can be used for the assessment of cell nuclear and nucleolar phenotype analysis methods. Cells are labeled with 3 different fluorophores: DAPI (4’,6-diamidino-2-phenylindole), a common stain for the nuclei, fibrillarin antibody (anti-fibrillarin) and ethidium bromide (EtBr), both used for nucleoli staining. In the dataset original images are in 1, 024 × 1, 024 × *Z* lattice (*Z* = {30, 50}). Every sub-volume is labeled as c0, c1, c2, representing the DAPI, anti-fibrillarin, and EtBr channels, respectively, **Fig. 1**. Accompanying meta-data are extracted from the original data. Binary masks are obtained by segmentation of the original data in c0 and c2 channels, see details in [5].

**Fig. 1.**
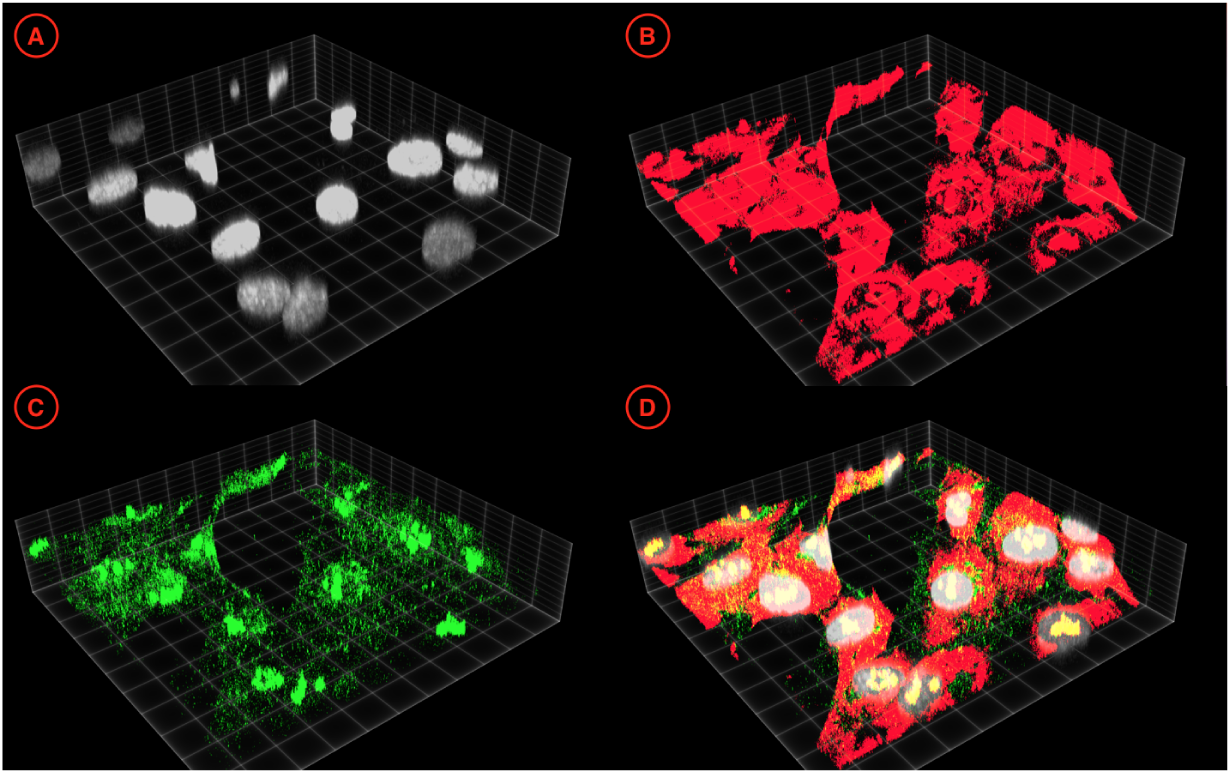
An exemplar 3D visualization of a data sub-volume from the fibroblast cell collection: (A) DAPI channel; (B) EtBr channel; (C) anti-fibrillarin channel; (D) a composite image. Adapted from [5] under a CC-BY 4.0 International license.

In this work we focus on images of primary human fibroblast cells. A part of this collection was subjected to a G0/G1 Serum Starvation Protocol used for cell cycle synchronization, has previously been shown to alter nuclear organization and to be reflected in changes in nuclear size and shape [8]. As a result, it contains 178 3D volumetric images of cells in the following phenotypic classes: (1) 64 sub-volumes of proliferating fibroblasts (PROLIF), and (2) 112 sub-volumes of the cell cycle synchronized by the serum-starvation protocol cells (SS). These classes serve as two categories in a binary morphology classification setting.

### 2.2 Data preprocessing

Fluorescent labels are not always specific to the object of interest and often produce noisy background (**Fig. 1**). In order to assess changes in the nuclear architecture, we first apply nuclear masks provided with the dataset to all 3 channels of original microscopy data. Due to the anisotropy in original data, we then re-scale volumes in *Z* dimension by a factor extracted from the corresponding meta-data. Since each of 1, 024 × 1, 024 × *Z* sub-volumes typically contains between 1 and 5 nuclei, we crop re-scaled volumes into smaller 256 × 256 × 57 sub-volumes, centered at the centroid of the corresponding nuclear mask and zero-pad them, when necessary. Finally, we produce 2D representation of sub-volumes by a maximum intensity projection along the *Z* dimension (**Fig. 2**). As a result, we create a set of 999 256 × 256 images per channel.

**Fig. 2.**
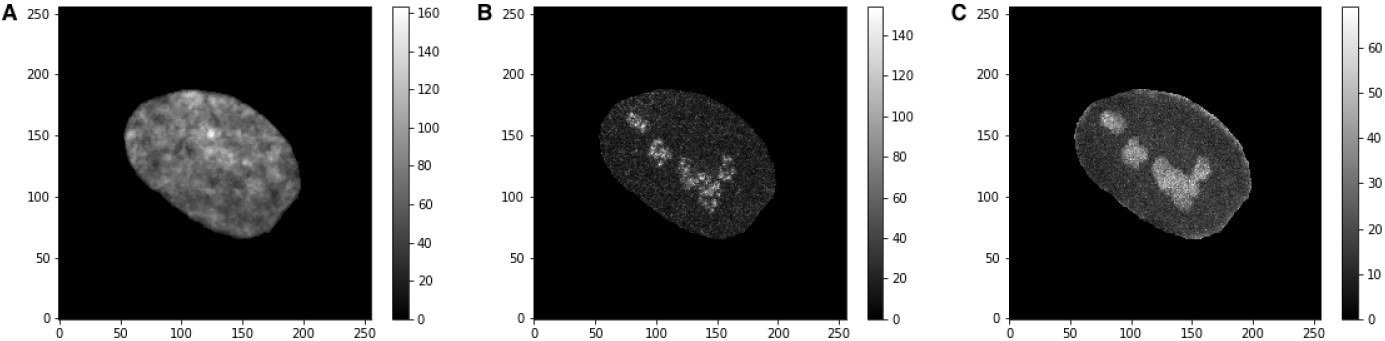
An exemplar visualization of 256 × 256 2D maximum intensity projections of a masked, re-scaled, and cropped fibroblast sub-volumes in: (A) DAPI channel, c0; (B) anti-fibrillarin channel, c1; (C) EtBr channel, c2.

### 2.3 Classification

We compare classification algorithms from scikit-learn, a popular Python machine learning toolkit [7], including Gaussian Naive Bayes (NB), Linear Discriminant Analysis (LDA), k nearest neighbors classifier (kNN), support vector machines with linear (SVM) and Gaussian kernels (RBF), Random Forest (RF), Extremely Randomized Trees (ET), and Gradient Boosting (GBM). All classifiers use default hyper-parameters. Every image is flattened into a 1D feature vector. Feature preprocessing includes subtracting the mean and scaling to unit variance of the training set. We assign the label of the whole image to every cell extracted from it. In order to assess batch effects in the intensity images and binary masks, we compare k-fold cross-validation (CV) scheme with the Leave-2-Opposite-Groups-Out (L2OGO) scheme, suggested in [5]. L2OGO ensures that: (1) all masks derived from one image fall either in the training or testing set, and (2) testing set always contains masks from 2 images of different classes.

## 3 Results

First, we evaluate the performance of algorithms for fibroblast nuclear classification using only 2D morphological information, i.e. binary masks. We compute AUROC per chennel using 2 different CV schemes: 20 splits in L2OGO and a 10 times repeated 4-fold CV. Results in **Table 1** do not show any apparent batch effects in the 2D classification setting in any of the channels, as performance levels L2OGO are only slightly lower compared to 4-fold CV. As expected, classifiers are not able to pick up complex morphological relationships from flattened binary vectors, even when 3 channels are combined. All such results were dominated by the classification performance using morphometry features manually extracted from binary masks, as described in [5]. The best overall result with L2OGO is achieved by the Gaussian SVM (RBF) classifier in with *AUROC* = 0.772±0.041, *AUPR* = 0.731 ± 0.063, and *F*1 = 0.682 ± 0.060.

Next, we evaluate the performance using only 2D pixel intensity information. Results in **Table 2** demonstrate a possibility of batch effects. The performance on the nuclear c0 channel does not benefit from the presence of additional information compared to only 2D masks. But nucleolar-stained channels c1 and c2 demonstrate 20% gain in performance even using more conservative L2OGO CV. However, L2OGO here leads to a large variance of the performance metric. On average, the EtBr channel (c2) seems to provide a sightly better representation of nucleolar structure comared to the anti-fibrillarin (c1). Almost all classifiers in both channels show results superior of those obtained with morphometric features, see **Table 1**. Combining all 3 channels gives the best result, demonstrating the complement nature of stains. The best overall result is achieved by the Gaussian SVM (RBF) classifier with *AUROC* = 0.990 ± 0.029, *AUPR* = 0.980 ± 0.040, and *F*1 = 0.877 ± 0.177).

## 4 Discussion

In order to establish baseline evaluation of simple pixel-based nuclear structure classification methods, we provide a comparison of a number of widely used machine learning algorithms on both binary and intensity 2D projections of 3D microscopic images. Although DAPI structure classification did not benefit from using the intensity information, our results indicate usefulness of intensities of nucleolar labels: anti-fibrillarin and EtBr. Nuclear morphometry extracted from binary masks seems to reflect most of the relevant changes. Increased potential for batch effects is only observed in classification of nucleolar structures in channels c1 and c2. Interestingly, combining 3 channels together seems to alleviate this issue and lead to near-perfect performance in L2OGO scheme.

**Table 1.** Classification AUROC (*mean ± std*) on binary masks for 2 cross-validation schemes (CV: 4-fold and L2OGO) and a number of algorithms (Clf: NB, LDA, kNN, SVM, RBF, RF, ET, GBM) per image channel (c0, c1, c2, and all 3 channels combined). For the reference, the last column contains performance measures obtained using voxel-based morphometry features extracted from binary masks [5].

| CV | Clf | Binary images |  |  |  | Morphology [5] |
| --- | --- | --- | --- | --- | --- | --- |
|  |  | c0 | c1 | c2 | c0c1c2 | c0c2 |
| 4-fold | kNN | 0.604 ± 0.056 | 0.615 ± 0.059 | 0.610 ± 0.059 | 0.620 ± 0.062 | 0.706 ± 0.030 |
| | SVM | 0.597 ± 0.067 | 0.583 ± 0.075 | 0.582 ± 0.054 | 0.650 ± 0.071 | 0.646 ± 0.090 |
| | RBF | 0.647 ± 0.053 | 0.648 ± 0.078 | **0.644 ± 0.056** | 0.678 ± 0.062 | **0.785 ± 0.027** |
| | RF | 0.635 ± 0.061 | 0.642 ± 0.061 | 0.635 ± 0.054 | 0.661 ± 0.064 | 0.735 ± 0.029 |
| | ET | 0.581 ± 0.059 | 0.577 ± 0.077 | 0.573 ± 0.058 | 0.651 ± 0.054 | 0.732 ± 0.034 |
| | GBM | **0.659 ± 0.049** | **0.658 ± 0.062** | 0.641 ± 0.061 | **0.721 ± 0.054** | 0.783 ± 0.032 |
| L2OGO | kNN | 0.582 ± 0.047 | 0.582 ± 0.047 | 0.582 ± 0.047 | 0.582 ± 0.047 | 0.695 ± 0.030 |
| | SVM | 0.557 ± 0.057 | 0.568 ± 0.050 | 0.555 ± 0.046 | 0.564 ± 0.053 | 0.650 ± 0.077 |
| | RBF | 0.616 ± 0.060 | 0.616 ± 0.060 | 0.616 ± 0.060 | 0.616 ± 0.060 | **0.772 ± 0.041** |
| | RF | 0.635 ± 0.038 | 0.636 ± 0.037 | 0.626 ± 0.050 | 0.627 ± 0.041 | 0.719 ± 0.039 |
| | ET | 0.637 ± 0.039 | 0.637 ± 0.039 | **0.641 ± 0.044** | 0.636 ± 0.042 | 0.717 ± 0.034 |
| | GBM | **0.642 ± 0.037** | **0.644 ± 0.038** | 0.639 ± 0.037 | **0.643 ± 0.041** | 0.758 ± 0.041 |

**Table 2.** Classification AUROC (*mean ± std*) on raw intensity images for 2 cross-validation schemes (CV: 4-fold and L2OGO) and a number of algorithms (Clf: NB, LDA, kNN, SVM, RBF, RF, ET, GBM) per image channel (c0, c1, c2, and all 3 channels combined).

| CV | Clf | Raw intensity images |  |  |  |
| --- | --- | --- | --- | --- | --- |
|  |  | c0 | c1 | c2 | c0c1c2 |
| 4-fold | kNN | 0.581 ± 0.059 | 0.771 ± 0.048 | 0.862 ± 0.041 | 0.865 ± 0.039 |
| | SVM | 0.610 ± 0.077 | 0.726 ± 0.080 | 0.829 ± 0.059 | 0.896 ± 0.043 |
| | RBF | 0.647 ± 0.058 | 0.814 ± 0.052 | 0.892 ± 0.0326 | 0.938 ± 0.026 |
| | RF | 0.630 ± 0.040 | 0.868 ± 0.039 | 0.890 ± 0.035 | 0.948 ± 0.022 |
| | ET | 0.606 ± 0.054 | 0.864 ± 0.045 | 0.875 ± 0.035 | 0.961 ± 0.021 |
|  | GBM | **0.673 ± 0.046** | **0.919 ± 0.031** | **0.912 ± 0.026** | **0.974 ± 0.011** |
| L2OGO | kNN | 0.552 ± 0.030 | 0.755 ± 0.207 | 0.826 ± 0.170 | 0.933 ± 0.044 |
| | SVM | 0.579 ± 0.053 | 0.671 ± 0.166 | 0.794 ± 0.183 | 0.964 ± 0.068 |
| | RBF | 0.579 ± 0.053 | 0.766 ± 0.261 | 0.844 ± 0.204 | **0.990 ± 0.021** |
| | RF | 0.579 ± 0.053 | 0.823 ± 0.202 | 0.841 ± 0.188 | 0.966 ± 0.057 |
| | ET | 0.613 ± 0.047 | 0.816 ± 0.209 | 0.839 ± 0.181 | 0.975 ± 0.034 |
| | GBM | **0.637 ± 0.045** | **0.844 ± 0.235** | **0.857 ± 0.204** | 0.990 ± 0.029 |

Presented evaluation has a number of drawbacks and requires further investigation. First, we only use flattened vectors of pixels, while there exist multiple methods for texture feature extraction, which may speed up the calculation. Alternatively, deep learning-based methods can be used for automatic feature learning [1, 6]. Second, we only evaluate performance on 2D maximum intensity projections of 3D images. Bigger study could further address similar issues in the original 3D space. Finally, we assume each nucleus in the same image to be representative of the phenotypic label that is provided for the whole image. This can be addressed by using methods that are robust to label noise [4].

